# Engineering the Marine *Pseudoalteromonas haloplanktis* TAC125 via pMEGA Plasmid Targeted Curing Using PTasRNA Technology

**DOI:** 10.1101/2024.12.13.628325

**Authors:** Angelica Severino, Concetta Lauro, Marzia Calvanese, Christopher Riccardi, Andrea Colarusso, Marco Fondi, Ermenegilda Parrilli, Maria Luisa Tutino

**Affiliations:** Dipartimento di Scienze Chimiche, Università degli Studi di Napoli Federico II, Complesso Universitario Monte S. Angelo, Via Cintia 4, 80126 Napoli, Italia; Dipartimento di Biologia, Università degli Studi di Firenze, Via Madonna del Piano 6, Sesto Fiorentino, Italia; Department of Integrative Structural and Computational Biology, The Scripps Research Institute, La Jolla, CA, 92037, USA; Istituto Nazionale Biostrutture e Biosistemi I.N.B.B., Via dei Carpegna, 19 00165 Roma, Italia

**Keywords:** *Pseudoalteromonas haloplanktis* TAC125, cold-adapted bacteria, pMEGA, megaplasmid, strain engineering, plasmid curing, PTasRNA gene silencing

## Abstract

Marine bacteria that have adapted to thrive in extreme environments, such as *Pseudoalteromonas haloplanktis* TAC125 (*Ph*TAC125), offer a unique biotechnological potential. The discovery of an endogenous megaplasmid (pMEGA) raised questions about its metabolic impact and functional role in this strain. This study aimed at streamlining the host genetic background by curing *Ph*TAC125 from the pMEGA plasmid using a sequential genetic approach. We combined homologous recombination by exploiting a suicide vector with the PTasRNA gene silencing technology to interfere with pMEGA replication machinery. This approach led to the construction of the novel *Ph*TAC125 KrPL^2^ strain, cured from the pMEGA plasmid, which exhibited no significant differences in the growth behaviour, though showcasing enhanced resistance to oxidative stress and a reduced capability of biofilm formation. These findings represent a significant achievement for understanding of the role of pMEGA plasmid and for the biotechnological applications of *Ph*TAC125 in recombinant protein production. This opens up the possibility to exploit pMEGA valuable genetic elements and further advancing the genetic tools for *Ph*TAC125.

## 1. Introduction

Much of life on Earth has evolved to thrive in cold environments, which constitute the most widespread habitat on the surface of our planet [1]. Despite the inherent challenges associated with surviving in these cold areas, psychrophilic (or psychrotolerant, or cold-adapted) microorganisms — including bacteria, archaea, yeasts, unicellular algae, and fungi — have remarkably adapted to life in these harsh conditions [2]. These micro-organisms can survive at temperatures below 7 °C, with optimal growth typically occurring at higher temperatures ranging from 20 to 30 °C [3].

Studies of bacterial communities in such environments have shown the widespread distribution and abundance of *gamma-proteobacteria*, which predominantly colonize both deep-sea and surface waters, as well as sea-costal sediments [4]. Among *gamma-proteobacteria*, the genus *Pseudoalteromonas* is particularly noteworthy because of its extensive distribution in marine environments where it represents a significant portion of the ocean bacterial community (2–3% in the surface ocean and 14% in the deep sea) [5]. Additionally, members of the *Pseudoalteromonas* genus are renowned for their exceptional environmental adaptability, enabling them to thrive in these extreme habitats [6,7].

Psychrophilic microorganisms counteract the deleterious effects of low temperatures by developing specific strategies in the form of finely tuned structural changes at the level of, for example, their membranes, constitutive proteins, and enzymes which explain their broad environmental adaptability [2,7]. In this context, *Pseudoalteromonas haloplanktis* TAC125 (*Ph*TAC125) is amongst the most valuable marine microorganisms, and it is considered a model organism of cold-adapted bacteria reaching high cellular density at the optimal temperature of 15 °C [8,9]. The fully sequenced and annotated multipartite genome of *Ph*TAC125 (composed of two chromosomes and pMtBL, a cryptic plasmid) [6,10], paved the way for understanding cold adaptation strategies in this bug, which in turn resulted to be: a) source of valuable bioactive compounds, such as anti-biofilm molecules [11,12] and antimicrobials [13,14] b) a tool for bioremediation [15,16] and c) as an exceptional host for recombinant production of “difficult proteins” preventing insoluble protein aggregates [17–20]. In this last regard, several expression vectors, based on the pMtBL replication signals and with either constitutive or inducible promoters [21,22], were developed to enable the recombinant production of many proteins both in planktonic [9,21] or biofilm cultures [23]. Additionally, different genetic tools were designed for *Ph*TAC125 including (a) a strategy for the construction of insertion/deletion genomic mutants exploiting homologous recombination, in which Giuliani and coworkers elaborated a technique for allelic exchange and/or gene inactivation by in-frame deletion and the use of a counterselection marker harnessing a non-replicative plasmid [24], and, (b) a conditional gene silencing system by PTasRNAs technology developed by Lauro et al. to effectively downregulate chromosomal genes expression of *Ph*TAC125 and achieve high levels of gene silencing [25].

Interestingly, in 2019, the resequencing of the *Ph*TAC125 genome using third-generation sequencing technologies uncovered a novel large plasmid, pMEGA, which is 64,758 bp in size and contains 52 open reading frames (ORFs) [10]. pMEGA shows limited similarity to *P. haloplanktis* chromosomes, but it shares a substantial nucleotide similarity with plasmids found in other marine bacteria, *Pseudoalteromonas arctica* and *Pseudoalteromonas nigrifaciens*, suggesting a unique evolutionary origin and potential adaptation to cold marine environments [10]. pMEGA is a low-copy number plasmid and it is considered a nonconjugative one. It encodes essential replication and stability functions, such as the RepB replication initiator protein, and the ParA and ParB proteins involved in partitioning and stability. Qi et al. leveraged the high nucleotide sequence conservation between the pMEGA plasmid and those from *P. arctica* and *P. nigrifaciens* to highlight similarities in the replication and partitioning functions, hypothesizing a Rolling Circle Replication (RCR) mechanism with a common regulation system [10]. pMEGA encompasses two types II toxin-antitoxin systems, the HipBA and the hybrid yefM-ParE [26]; it harbours genes coding for defence mechanisms against bacteriophages, including type I and type IV restriction-modification systems as well as several proteins with a role in cell metabolism (TonB-dependent receptors, an aminotransferase, a nitronate monooxygenase, an epimerase, and an acetyltransferase).

More recently, pangenome studies revealed that multipartite genomes are widely distributed in the class *gamma-proteobacteria* in which the major representatives are *Vibrionaceae* [27] and *Pseudoalteromadaceae* (prevalently *Pseudoalteromonas spp*.) [28]. Using a phylogenetic approach and timescale analysis on *Pseudoalteromonas*, Liao et al. (2019) showed that chromosomes and chromids (carrying essential housekeeping genes localized on the chromosome in other species) have always coexisted and that chromids apparently originated from a megaplasmid that over time obtained essential genes enhancing the hypothesis that megaplasmids are commonly distributed in this genus [6,29]. However, some megaplasmids, like pMEGA that lack their own conjugative machinery either rely on the conjugative systems of other plasmids for transmission (i.e., they are mobilizable) or may not be mobile at all. In the long run, these plasmids may either impose a metabolic burden and evolve into efficient selfish mobile elements or become mutualists, dependent on the host for replication and transmission [30]. Loss or removal of the megaplasmid, might help highlight other aspect of the physiological and functional roles of that conserved megaplasmid in *Ph*TAC125.

In this study, we explored the genomic and transcriptomic landscape of the pMEGA plasmid to build up a finely tuned strategy tailored for its curing. We developed a sequential genetic scheme through the combination of two different genetic methodologies previously developed in *Ph*TAC125. We first edited pMEGA by integrating the pAT suicide vector carrying a selection marker, necessary to distinguish wild type from cured bacteria. Then we used the PTasRNA gene silencing technology to target and interfere with the replication machinery of the plasmid. We successfully removed the pMEGA plasmid from the bacterium which resulted in the construction of a novel strain named KrPL^2^. The phenotypical characterization showed that this strain exhibited comparable growth dynamics to its progenitor strain, demonstrating that the removal of pMEGA did not significantly impact the strain growth or survival across different temperatures and media conditions. Additionally, plasmids that share similarities in their replication or partitioning systems, cannot stably coexist in the same bacterial cell leading to plasmid incompatibility [31]. Hence, further evaluations of plasmid incompatibility, stability, and copy number revealed that, despite the removal of pMEGA, segregation incompatibility of *rep*-based plasmids persisted, indicating the involvement of additional mechanisms beyond the replication associated genes. This study also confirmed that the removal of pMEGA did not affect pMtBL-derived plasmid stability or copy number under the conditions tested. Also, this work demonstrates the efficacy of PTasRNA technology in curing plasmids with rolling-circle replication, extending the application of this methodology for future applications in genetic manipulation and functional studies in psychrophilic bacteria.

## 2. Materials and Methods

### 2.1 Bacterial Strains, Media and Plasmids

*E. coli* TOP10 (*mcrA*, Δ(*mrr-hsdRMS-mcrBC*), *ϕ80lacZ* (del) *M15, ΔlacX74, deoR, recA1, araD139*, Δ(*ara-leu*) - 7697, *galU, galK, rpsL* (SmR), *endA1, nupG*) was used as host for cloning procedures. *E. coli* strain S17-1(λpir) [*thi, pro, hsd* (*r*− *m+*) *recA*::RP4-2TCr::Mu Kmr::Tn7 Tpr Smr λpir] [21,32] was used as a donor in intergenic conjugation experiments. *Pseudoalteromoans haloplanktis* TAC125 (*Ph*TAC125) KrPL, a strain cured from the endogenous pMtBL plasmid [21], was used as a host for the homologous recombination experiment. *Ph*TAC125 KrPL *ins*DNApolV mutant was used as host for the expression of the anti-sense RNA in the conditional gene silencing protocol. *Psychrobacter sp*. TAD1 (NCBI:txid81861) was used as a control of the incompatibility assay.

*E. coli* strains were grown in LB broth (10 g/L bacto-tryptone, 5 g/L yeast extract, 10 g/L NaCl) at 37 °C and the recombinant strains were treated with either 34 μg/mL chloramphenicol or 100 μg/mL ampicillin, depending on the selection marker of the vector. *Ph*TAC125 strains were cultured in TYP medium (16g/L bacto-tryptone, 16 g/L yeast extract, 10 g/L NaCl), GG 10-10 medium (10 g/L L-glutamic acid monosodium salt monohydrate, 10 g/L D-gluconic acid sodium salt, 10 g/L NaCl, 1 g/L NH_4_NO_3_) [9] for recombinant production of PTasRNA*repB* and GG 5-5 (5 g/L D-gluconate, 5 g/L L-glutamate, 10 g/L NaCl, 1 g/L NH_4_NO_3_) accordingly to the experimental set up. During the experiments, GG 10-10 and GG 5-5 are supplemented with salts Schatz (1 g/L K_2_HPO_4_, 200 mg/L MgSO_4_·7H_2_O, 5 mg/L FeSO_4_·7H_2_O, 5 mg/L CaCl_2_) in sterile conditions.

When necessary, chloramphenicol was added to solid and liquid media at 12.5 μg/mL and 25 μg/mL concentrations, respectively. Ampicillin was always used with a concentration of 100 μg/mL, instead.

Competent cells of *E. coli* strains were made using the calcium chloride method and transformed with the appropriate vector through heat-shock transformation method [33]. *Ph*TAC125 cells were transformed with the appropriate vector via intergeneric conjugation as reported in Tutino et al. 2001 [32].

The recombinant plasmids used in this work are listed in Table 1. pMAV was used during the plasmid copy number assessment and the construction is reported in Sannino et al., 2017 [9]. pKT240 and pUC-oriT-pTAUp plasmids along with pMAV plasmid were employed during the incompatibility assay. The pAT-eGFP was used for the construction of the pAT-VS-HR*umuC* which was designed for the pMEGA curing strategy. The pB40-79-PTasRNA*lon*, pB40-79C-PTasRNA*lon*, and pB40-79-PTasRNA*repB* were designed and used in the PT-antisense interference experiment for the selection of the pMEGA cured clones. The pB40-79-PTasRNA*rep*B was also used for the heterologous plasmid stability assessment.

**Table 1.**
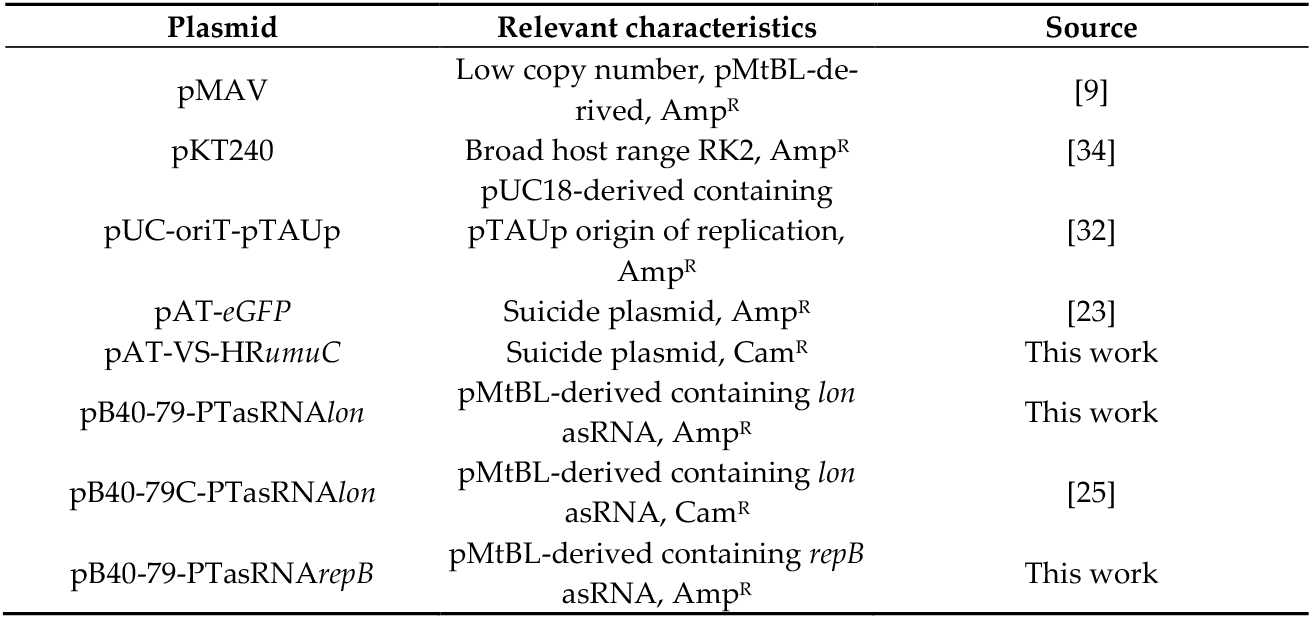
List of plasmids used in this study.

| Plasmid | Relevant characteristics | Source |
| --- | --- | --- |
| pMAV | Low copy number, pMtBL-derived, Amp<sup>R</sup> | [9] |
| pKT240 | Broad host range RK2, Amp<sup>R</sup> | [34] |
| pUC-oriT-pTAUp | pUC18-derived containing pTAUp origin of replication, Amp<sup>R</sup> | [32] |
| pAT-eGFP | Suicide plasmid, Amp<sup>R</sup> | [23] |
| pAT-VS-HRumuC | Suicide plasmid, Cam<sup>R</sup> | This work |
| pB40-79-PTasRNA<sub>lon</sub> | pMtBL-derived containing *lon* asRNA, Amp<sup>R</sup> | This work |
| pB40-79C-PTasRNA<sub>lon</sub> | pMtBL-derived containing *lon* asRNA, Cam<sup>R</sup> | [25] |
| pB40-79-PTasRNA<sub>repB</sub> | pMtBL-derived containing *repB* asRNA, Amp<sup>R</sup> | This work |

### 2.2 Bioinformatic Analyses and Transcriptomic Data Collection

To identify regions of similarity between pMEGA and other prokaryotic genomes an initial BlastN similarity search against the NCBI nr/nt nucleotide database was performed using the NIH web interface in its default parameters [35]. Similarly, BlastP similarity searches of the *Ph*TAC125 proteome were then used to identify pMEGA proteins homologous to *Pseudoalteromonas sp*. KG3 and *Pseudoalteromonas sp*. PS1M3 through the NCBI non-redundant protein sequences (nr) database.

For the PTasRNA interference assay, prediction of RNA secondary structures was performed using the mFold website with default settings [36]. Gene expression data for *Ph*TAC125 cultured at two temperature conditions (0°C and 15°C, respectively) were collected from Riccardi et al. [37], and the information relative to pMEGA transcriptome was kept for further analyses. Sequence annotations for pMEGA CDS and FASTA were downloaded from the NCBI, using accession number NZ_MN400773.1, last accessed on October 10^th^ 2024.

### 2.3 Plasmid Construction

The suicide vector pAT-VS-HR*umuC* was obtained through gateway cloning procedures. The HR*umuC* homology region of 259 bp located on pMEGA plasmid was PCR-amplified using the bacterial genomic sample of *Ph*TAC125 as DNA template, which includes the pMEGA plasmid, and a pair of oligos forward and reverse listed in Supplementary Table S3 (*umuC*_*Not*I fw, *umuC*_*Asc*I rv). The PCR fragment was purified and subjected to *Not*I/*Asc*I double digestion and cloned inside the pAT-eGFP vector previously hydrolyzed with the same restriction enzymes. The homology region sequence was inserted between the *oriC* and *oriT* of the pAT-eGFP vector, resulting in a plasmid lacking the psychrophilic origin of replication, thus yielding the pAT-VS-HR*umuC* suicide vector. The pB40-79-PTasRNA*lon* vector, a derivative of pB40-79C-PTasRNA*lon* used in our previous work [25], was constructed through a cut and paste procedure where both plasmids were double digested with *SphI*/*AscI* restriction enzymes. These cleavages generate a fragment A of 2,276 bp carrying the *ori*C and *amp*R regions, whereas fragment B of 2,815 bp delivers the Paired-Termini and antisense region, along with the *oriR, oriT*, MCS and *Ph*LacR and PLacZ promoter regions. Joining the two fragments led to the construction of the pB40-79-PTasRNA-*lon* vector in which the chloramphenicol selection marker has been replaced with the ampicillin resistance gene.

The construction of pB40-79-PTasRNA*rep*B was carried out by exploiting *Pst*I and *Xho*I restriction sites in pB40-79-PTasRNA*lon*. These sites were purposefully designed by Lauro et al. to directly exchange the antisense sequences without altering the paired termini sequences upstream and downstream of the antisense RNA. The asRNA*rep*B was PCR amplified (Phusion High-Fidelity DNA polymerase - Thermo Fischer Scientific) using pMEGA plasmid as a template and a pair of oligos forward and reverse listed in Supplementary Table S1 (*Xho*I-asRNA*rep*B fw, *Pst*I-asRNA*rep*B rv). The PCR fragments and the pB40-79-*Phlon* were double hydrolyzed with *Pst*I/*Xho*I restriction enzymes resulting in the pB40-79-PTasRNA*rep*B vector. The nucleotide sequence of the synthetic asRNA*repB* in this work is listed in Supplementary Table S2 Bacterial DNA kit (D3350-02, E.Z.N.A™, OMEGA bio-tek, Norcross, GA, USA) was used for DNA genomic extraction following the manufacturer’s instructions.

### 2.4 PTasRNA Interference

Gene silencing using the PTasRNA*rep*B was performed by culturing the bacteria in 20 mL GG 10-10 medium, as described in *2.1 Bacterial strains, growth conditions and bacterial manipulation*, supplemented with 100 μg/mL ampicillin in 100 mL Erlenmeyer flasks. Also, the non-recombinant Krpl *ins*DNApolV strain was used as a negative control of the experiment grown in non selective conditions.

Strains were streaked on TYP agar plate supplemented with 100 μg/mL ampicillin and incubated at 15 °C for three days. A single colony was inoculated in 3 mL TYP medium supplemented with 100 μg/mL ampicillin at 15 °C overnight in 25 mL inoculum glass tube. Cultures were diluted to 1:100 in 10 mL GG 10-10 supplemented with salts Schatz and 100 μg/mL ampicillin in 100 mL Erlenmeyer flask and incubated at 15 °C with shaking. Following, a 0.35 OD/mL dilution in 20 mL GG 10-10 medium supplemented with salts Schatz and 100 μg/mL ampicillin was performed and incubated at 15 °C for 8 hours in 100 mL Erlenmeyer flask with shaking.

The inoculum was performed at 0.1 OD/mL, and the induction was carried out with 10 mM IPTG when the cells reached a value of 1.5 OD/mL. Samples were collected at 8, 24 and 32 hours post-induction. 0.1 OD/mL (1-2 x 10^7^ CFU/mL) cultures were diluted 1:10^4^ and 100 μL were seeded on non-selective TYP agar. The incubation was performed at 15 °C for 3 days obtaining about 100-200 colonies. The grown colonies from each plate (8, 24 and 32 hours post-induction) were replicated on 70 mL TYP agar plate in the presence and absence of chloramphenicol as a selective agent and then incubated at 15 °C.

The clones that do not replicate on TYP agar supplemented with chloramphenicol but easily grow on TYP agar without the selective agent, express the PTasRNA*repB*. These clones are subjected to PCR colony screening. A single colony of *Ph*TAC125 was resuspended in 30 µL of deionized MilliQ water, boiled for 10 minutes and spun down for 10 minutes. The supernatant (1 µL) was used as DNA template for PCR reaction using Taq Polymerase (Thermo Scientific™).

### 2.5 Growth Cultures Conditions at 0 °C, 15 °C and 20 °C

Each strain was cultured at 0 °C, 15 °C and 20 °C temperatures in TYP medium and GG 5-5 media supplemented with salts Schatz. The strains were streaked on TYP agar plate and incubated at 15 °C for three days.

A single colony was inoculated in 3 mL TYP medium at 15 °C overnight in 25 mL inoculum glass tube. Cultures at 15 °C were performed diluting in 10 mL TYP or GG 10-10 supplemented with salts Schatz in 100 mL Erlenmeyer flask and incubated at 15 °C with shaking. Following, a 0.35 OD/mL dilution in 20 mL TYP medium or GG 5-5 medium supplemented with salts Schatz was performed and incubated at 15 °C for 8 hours in 100 mL Erlenmeyer flask with shaking. The inoculum was performed at 0.1 OD/mL at 15 °C in 50 mL TYP or GG 5-5 medium supplemented with salts Schatz in 250 mL Erlenmeyer flask with shaking. Points of the growth curves were collected every 3 hours.

Cultures at 20 °C and 0 °C were carried out pre-adapting the pre-inoculum at the desired temperature to avoid drastic changes in growth conditions. The experimental steps were carried out following the 15 °C cultures set up adding a pre-adaptation step at 20 °C or 0 °C before the inoculum. The 20 °C growth curves were carried out for 72 hours and points were collected every 2 hours, whereas the 0 °C curves were carried out for about 200 hours and points were collected every 8 hours. The experiment was set up as biological duplicate in which the error is reported as standard deviation determined by Excel *dev.st.c* function.

A significance level of 0.05 was chosen prior to the experiments. Two-sided Student’s *t*-tests of two samples were conducted using *GraphPad Prism 8* to test the null hypothesis that there is no difference between means.

### 2.6 Segregation Stability and Incompatibility Assay

Plasmid segregation stability was assayed under non-selective conditions for over 150 generations. Each strain was selected from a single colony on TYP agar supplemented with 100 μg/mL ampicillin and grown in 3 mL TYP medium at 15 °C for 24 hours with shaking. Then, a dilution was performed in 3 mL GG 5-5 medium supplemented with salts Schatz and 100 μg/mL ampicillin, and incubated for 24 hours at 15 °C with shaking. The stability assay was performed by growing the bacteria at 15 °C in 5 mL GG 5-5 medium in non-selective condition inoculating at 0.05 OD until the exponential phase (1-2 OD). This step was performed daily until the desired number of generations.

The percentage of recombinant cells was assessed by seeding diluted samples on non-selective TYP agar. 0.1 OD/mL (1-2 x 10^7^ CFU/mL) culture was diluted 1:10^4^ and 100 μL were seeded, obtaining about 100-200 colonies after incubation at 15 °C for three days. 30-33 colonies were taken from each plate and replicated on both selective (100 μg/mL ampicillin) and non-selective plates which were incubated at 15 °C for 24 hours. Each culture was performed as a biological duplicate and the average number of colonies resistant to 100 μg/mL ampicillin was calculated. Plasmid rate loss was calculated by the number of colonies grown on the TYP-agar plates with ampicillin divided by the number of colonies grown on the TYP-agar plates without ampicillin.

During the incompatibility assay, the *rep*-based replication plasmids were transferred to *Ph*TAC125 strains via intergeneric conjugation using *E. coli* strain S17-1(λpir) as donor strain. pMAV plasmid represents a positive control of the intergeneric conjugation experiment. *Psychrobacter sp*. TAD1 represents a positive control for pUC-oriT-pTAUp plasmids intergeneric conjugation. Selection of trans-conjugative clones was performed at the temperature of 4 °C using 100 μg/mL ampicillin. The clones were selected and subjected to replica plating using 100 μg/mL ampicillin.

### 2.7 Plasmid Copy Number Quantification

Relative plasmid copy number (PCN) was determined by the quantitative PCR (qPCR) technique. Amplification and data analysis were performed with a StepOne Realtime PCR System (Applied Biosystems, Foster City, CA, USA) along with the SYBR® Green PCR Kit (Applied Biosystems, Foster City, CA, USA). The gene with locus tag *PSHA_RS10135* (alternatively known as *PSHAa2051*) was employed to detect chromosomal DNA in the samples, while the *AmpR* gene was used for the detection of the plasmid. Each couple of primers (prom 7 fw, prom7 rv, BlaM fw, BlaM rv, Supplementary Table S1 was selected using the free Primer 3 web software.

For PCN estimation of unknown samples, the total DNA was extracted from 1 OD_600_ cell pellets of *Ph*TAC125 exponential phase cultures grown in both TYP and GG 5-5 media. DNA extraction was performed using the Bacterial DNA kit (D3350-02, E.Z.N.A™, OMEGA bio-tek, Norcross, GA, USA) following the manufacturer’s instructions. DNA concentrations were measured with a NanoDrop TM 1000 Sp spectrophotometer (Thermo Fisher Scientific, Waltham, MA, USA) at the absorption of 260 nm and purity was assessed by measuring the A260/280 and A260/230 ratios of the extracted DNA. PCR reactions were prepared in 10 μL mixtures containing 1 × PowerUp SYBR Green Master Mix (Applied Biosystems, Foster City, CA, USA) with ROX as passive reference dye and Uracil-DNA glycosidase (UDG) to eliminate contaminations, 400 nM of each primer and 1 μL of sample and the reaction master mixes were aliquoted in three wells of a reaction plate to perform a technical replicate. Finally, the plate was sealed with an adhesive cover (Applied Biosystems, Foster City, CA, USA).

The thermal cycling protocol was as follows: UDG activation for 2 min at 50 °C; initial denaturation for 10 min at 95 °C; 40 cycles of denaturation for 15 s at 95 °C alternated with annealing/extension steps for 1 min at 60 °C. Each reaction was performed in triplicate. Standard curves were developed using 10-fold serial dilutions of a random real sample (from 6 × 10^3^ to 6 pg of total DNA). In each dilution series, either the chromosomal gene or the plasmid gene was the target.

The amplification efficiency (E) of each gene was calculated from the slope of the relative standard curve (E = 10(−1/slope)). The relative PCN was estimated with the following equation: = (E_c_)^Ctc^/(E_p_)^Ctp^, considering the Ct values for the two amplicons (chromosome-c and plasmid-p) and the amplification efficiency of the plasmid (Ep) and chromosomal gene (Ec) [38].

### 2.8 H_2_O_2_ Disk-inhibition and Motility Assay

The disk-diffusion assay was performed in the presence of 265 mM H_2_O_2_. A single colony of each strain was inoculated in 3 mL of TYP media in non-selective conditions. Cells at the stationary phase were diluted to 0.2 OD/mL in 6.5 mL TYP soft agar (0.4% agar) and poured into a plate. Disks of Whatman filter paper (0.6 cm) were treated with 3 μL of 3.6% H_2_O_2_, placed in the centre of each plate, and incubated at 15 °C. After 24h of incubation, the disk of inhibition was measured as the cm of the diameter of inhibition. The experiment was performed as five independent replicates.

The motility assay was carried out for 72 hours of incubation. Each strain was streaked on TYP agar and incubated at 15 °C for three days. A single colony was inoculated in 3 mL TYP medium and incubated at 15 °C for 24 hours. Upon reaching the stationary phase, the cells were diluted at 2 OD/mL in TYP medium and 25 µL were spotted on sterile Whatman filter paper placed at the centre of a 0.3% soft agar TYP medium plate. After drying the spot in sterile conditions for 10 minutes, the plates were incubated at 15 °C. The length of the path (cm) was measured every 24 hours. The experiment was performed as three technical replicates.

### 2.9 Biofilm Formation Assay

Each strain was grown in TYP medium and GG 5-5 medium at 15 °C and 0 °C for 144 hours and points were collected every 24 hours. A single colony of each strain was inoculated in 2 mL of TYP at 15 °C for 24 hours. Cells were diluted to 0.35 OD/mL in 20 mL TYP media and GG 5-5 supplemented with salts Schatz. The pre-inoculum was split, and 10 mL were incubated at 15 °C for 8 hours with shaking in 100 mL Erlenmeyer flasks and the other 10 mL were incubated at 0 °C for 72 hours for cultures temperature preadaptation. For each temperature and media condition, the wells of a sterile 24-well flatbottomed polystyrene plate were filled with 1 mL of a medium with a 0.2 OD/mL dilution of the Antarctic bacterial culture in the exponential growth phase in static condition. The plates were incubated either at 15 °C or 0 °C and the kinetics of biofilm formation were carried out for 24-, 48-, 72-, 96-,120-, and 144-hours.

After rinsing with PBS, the adherent cells were stained with 0.1% (w/v) crystal violet, rinsed twice with double-distilled water, and thoroughly dried. Subsequently, the dye bound to the adherent cells was solubilized with 20% (v/v) acetone and 80% (v/v) ethanol. After 10 min of incubation at room temperature, the OD_590_ nm was measured to quantify the total biomass of biofilm formed in each well. The OD_590_ nm values reported were obtained by subtracting the OD_590_ value of the control obtained in the absence of bacteria. Each data point was composed of four independent samples.

### 2.10 Statistics and Reproducibility of Results

Data were statistically validated using the t-Student test comparing the mean values to the wt control. The significance of differences between the mean values was calculated using a two-tailed Student’s *t*-test and a p < 0.05 was considered significant. For multiple comparison, an ordinary one-way ANOVA test was performed using the Bonferroni test correction. A significance level of 0.05 was chosen prior to the experiments. Statistical analyses were conducted using *GraphPad Prism 8*.

## 3. Results

### 3.1 Genomic and Transcriptomic Profiling of the pMEGA Plasmid

In 2019, third-generation sequencing technologies led to the reannotation of the *Pseudoalteromonas haloplanktis* TAC125 genome, unveiling the presence of pMEGA. Previous NCBI nucleotide searches of this megaplasmid performed by Qi et al. with the NCBI [10] revealed only two significant hits to plasmids from *Pseudoalteromonas arctica* (36% of identity) and *Pseudoalteromonas nigrifaciens* (34% of identity), and scarce similarity to the chromosomes I (5%) and II (2%) of *Ph*TAC125 [10].

Currently (October 2024), pMEGA nucleotide similarity searches against the NCBI nucleotide collection database (nr/nt) revealed two more hits covering 33% and 26% of the pMEGA sequence, corresponding to endogenous plasmids of *Pseudoalteromonas sp*. KG3 and *Pseudoalteromonas sp*. PS1M3, respectively (Table 2).

**Table 2.** BlastN similarity searches against the NCBI nr/nt nucleotide database.

| Description | Query coverage | E-value | Identity (%) | Accession |
| --- | --- | --- | --- | --- |
| *Pseudoalteromonas* sp. KG3 plasmid pPsKG3-1 | 33% | 0.0 | 95.04 | CP098525.1 |
| *Pseudoalteromonas* sp. PS1M3 plasmid pPS1M3-2 | 26% | 0.0 | 95.05 | AP022626.1 |
| *Vibrio furnissii* strain 104486766 chromosome I | 8% | 0.0 | 87.44 | CP100422.1 |
| *Vibrio furnissii* strain VFN3 chromosome I | 8% | 0.0 | 87.44 | CP089603.1 |
| *Pectobacteriaceae* bacterium CE90 chromosome | 8% | 0.0 | 87.59 | CP129114.1 |

The former hit matches the 184,073 bp long pPsKG3_1 plasmid from *Pseudoalteromonas sp*. KG3 isolated from cheese rind (NCBI:txid2951137) showing 95.04% of identity with a region of pMEGA encoding for endonuclease subunit R and S (NDQ71_24070, NDQ71_24080) of type I restriction modification system, a gene encoding the toxin HipA (NDQ71_24235) and a gene encoding the HNH endonuclease (NDQ71_24225).

The latter hit matches plasmid pPS1M3-2 (NZ_AP022626), which is a large plasmid of 27,013 bp harboured by *Pseudoalteromonas sp*. PS1M3, a marine bacterium isolated from sea sediment from the Boso Peninsula in Japan [39]. This plasmid shares 95.05% identity with a region of pMEGA corresponding to *parA* and *parB* genes (98% of identity taken individually), encoding the plasmid partitioning system, and *hsdR*_2 gene (95% of identity) encoding a type I restriction endonuclease. Overall, the NCBI nucleotide BLAST analysis results showed hits against *Pseudoalteromonas* species, spanning from marine bacteria (i.e., *P. arctica* and *P. nigrifaciens*) to non-marine bacteria (i.e., *Pseudoalteromonas sp. KG3*). However, there were also significant matches with *Vibrio* species (CP100422.1, *Vibrio furnissii* chromosome I; CP089603.1, *Vibrio furnissii* strain VFN3 chromosome I), with 8% query coverage and 87.44% identity, particularly in intergenic regions and pseudogenes (Table 2).

Protein similarity searches of pMEGA against the NCBI non-redundant protein sequences (nr) revealed that pMEGA-encoded proteins show similarity with two proteins of *Pseudoalteromonas sp*. KG3 and *Pseudoalteromonas sp*. PS1M3. One hit corresponds to the ParA protein (WP_086998901, WP_024606458) sharing more than 99% of identity, while the other is an AAA family ATPase protein (WP_301562706, WP_006793244) sharing more than 30% of identity (Table 3).

**Table 3.** Significant results of BlastP similarity searches of the proteins residing on pMEGA against the nr database.

| Description | Protein annotation | Query coverage | E-value | Identity (%) | Accession |
| --- | --- | --- | --- | --- | --- |
| *Pseudoalteromonas* sp. KG3<br>plasmid pPsKG3-1 | ParA family protein | 100% | 0.0 | 99.75 | WP_086998901 |
|  | AAA family ATPase protein | 89% | 2,00E-53 | 32.61 | WP_301562706 |
| *Pseudoalteromonas* sp. PS1M3<br>plasmid pPS1M3-2 | ParA family protein | 100% | 0.0 | 99.75 | WP_024606458 |
|  | AAA family ATPase protein | 91% | 5,00E-55 | 32.36 | WP_006793244 |

Additionally, the 64,789 bp sequence of the pMEGA plasmid encompasses 52 annotated open reading frames (ORFs) classified into 6 different functional categories [10]. Following up on differential gene expression analyses that we conducted in a previous work for the same bacterium under two temperature conditions [37], we re-examined the transcriptional profile focusing specifically on the genes harboured by pMEGA. Among the 52 encoding genes, we identified 27 differentially expressed genes of which 2 genes and 25 genes are respectively down- and up-regulated at 15 °C (Table 4).

**Table 4.**
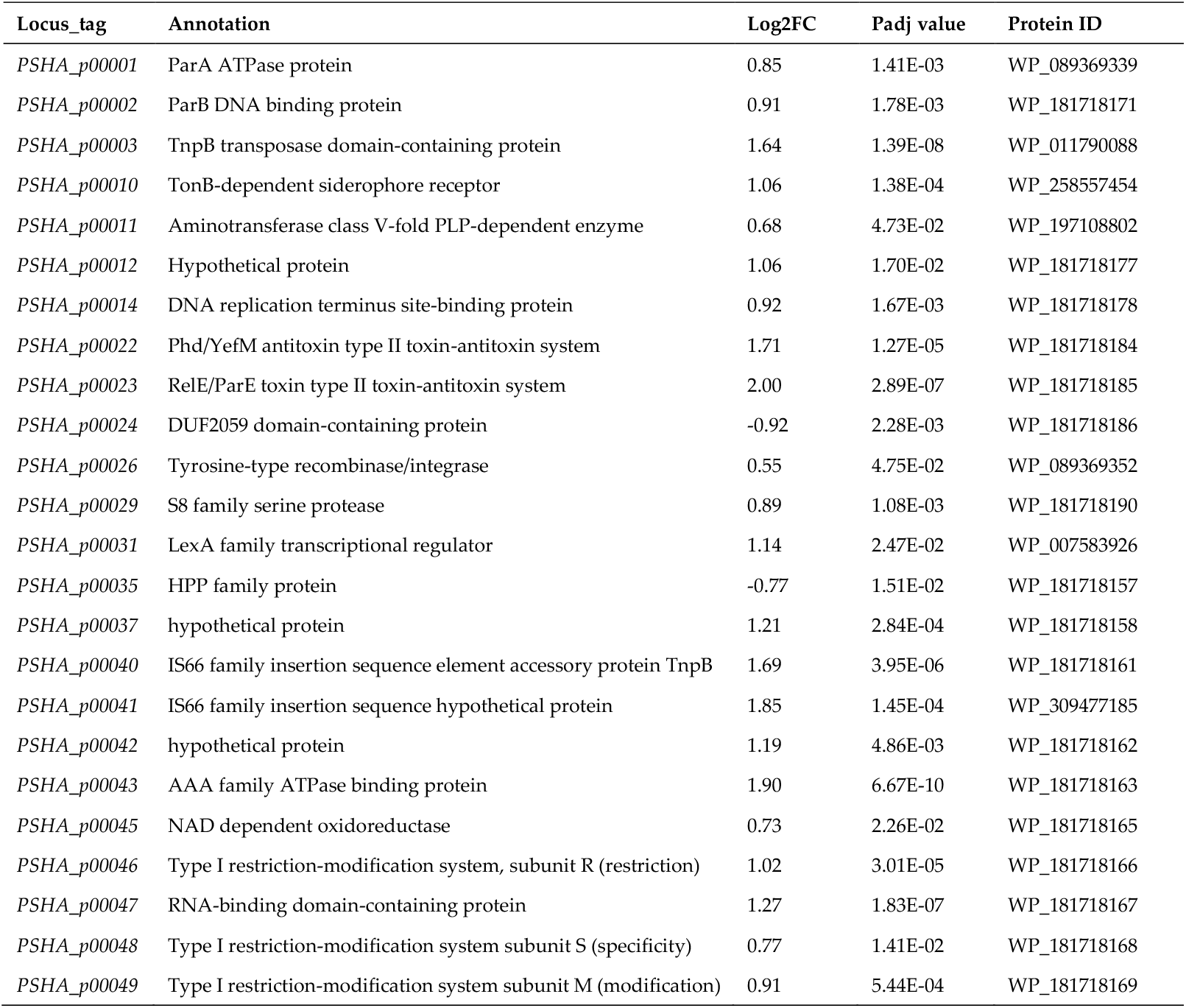
pMEGA differentially expressed genes (DEGs) at 15 °C and 0 °C.

The two genes (*PSHA_p00024, PSHA_p00035*) down-regulated at 15 °C are annotated as a DUF2059 domain-containing protein (PF09832) and HPP protein (PF04982) respectively but their function is unknown or uncertain. The uncharacterized DUF2059 is a domain predominantly found in bacterial proteins. The HPP is an integral membrane protein, equipped with four transmembrane spanning helices, that might function as a transmembrane transporter.

On the other hand, within the 25 genes upregulated at 15 °C, three are annotated as hypothetical proteins (*PSHA_p00012, PSHA_p00037, PSHA_p00042*). Moreover, we identified six genes with higher upregulation at 15 °C. Two of them, *PSHA_p00001* and *PSHA_p00002* genes encoding the ParA and ParB proteins respectively (PF18607, PF08775), regulate plasmid segregation system of pMEGA. Notably, among the top five identified genes *PSHA_p00022* and *PSHA_p00023* show the highest rate of upregulation at 15 °C, 5-fold compared to 0 °C.

These genes represent the hybrid toxin-antitoxin systems encoding the YefM antitoxin (IPR036165) and the ParE toxin (IPR007712), respectively, involved in pMEGA maintenance and stability [10]. Taking this into consideration, the upregulation of *par* genes and the toxin-antitoxin modules at 15 °C, the optimal growth temperature for the bacterium, might reflect the higher cellular replication rates and hence the increased demand for an efficient and robust partitioning system to ensure plasmid distribution to the daughter cells. Interestingly, the last gene (*PSHA_p00043*) encodes an AAA+ protein or ATPase which exerts diverse activities such as molecular chaperone, ATPase subunits of protease, helicase, translocation of macromolecules or nucleic-acid stimulated ATPases (PF13191). This protein might be involved in different stages of cellular growth, such as nutrients uptake, DNA replication and repair. Besides the identified differentially expressed genes (DEGs), it is worth noting the *PSHA_p00052* gene is annotated as a replication initiation protein (RepB) (PF01051). The transcriptional level of the gene is not down or upregulated at the analyzed temperatures. This is expected considering that previous studies indicate the product of *repB* gene as a key player in the rolling-circle mechanism, coding for the plasmid replication initiator, RepB, crucially ruling the duplication of pMEGA [10]. Considering that *repB* is an essential gene for plasmid maintenance, it stands out as a good candidate for plasmid curing.

### 3.2 Design of the Curing Strategy

Although pMEGA is endowed with several genes, none can be considered a useful selection marker to track down plasmid depletion. Thus, to achieve proper curing of pMEGA, a sequential genetic scheme was developed as follows: a) gene inactivation by in-frame insertion through homologous recombination to insert an antibiotic selection marker, and b) gene silencing of plasmid replication elements by PTasRNA technology.

The first step was designed to introduce the chloramphenicol selection marker within pMEGA, developing a novel mutant strain of *Ph*TAC125, named KrPL insPolV. Construction of the KrPL insPolV mutant was carried out by exploiting the pAT non-replicating plasmid (suicide vector, pAT-VS-HR*umuC*) to introduce the chloramphenicol resistance gene (*cmR*) into pMEGA. Also, the suicide vector comprises the 259 bp homology region of the *umuC* gene (HR*umuC*) which is a fragment of the *PSHA_00030* gene located on pMEGA (named A in Figure 1a) and the gene of the eGFP under the *placZ* promoter (optimized for *Ph*TAC125). Homology recombination requires the mobilization of the suicide vector pAT-VS-HR*umuC* into the host cell *Ph*TAC125 KrPL, a strain devoid of the pMtBL plasmid [32], by intergeneric conjugation harnessing the *E. coli S17-1 λpir* strain [21,25,32] as the donor.

**Figure 1.**
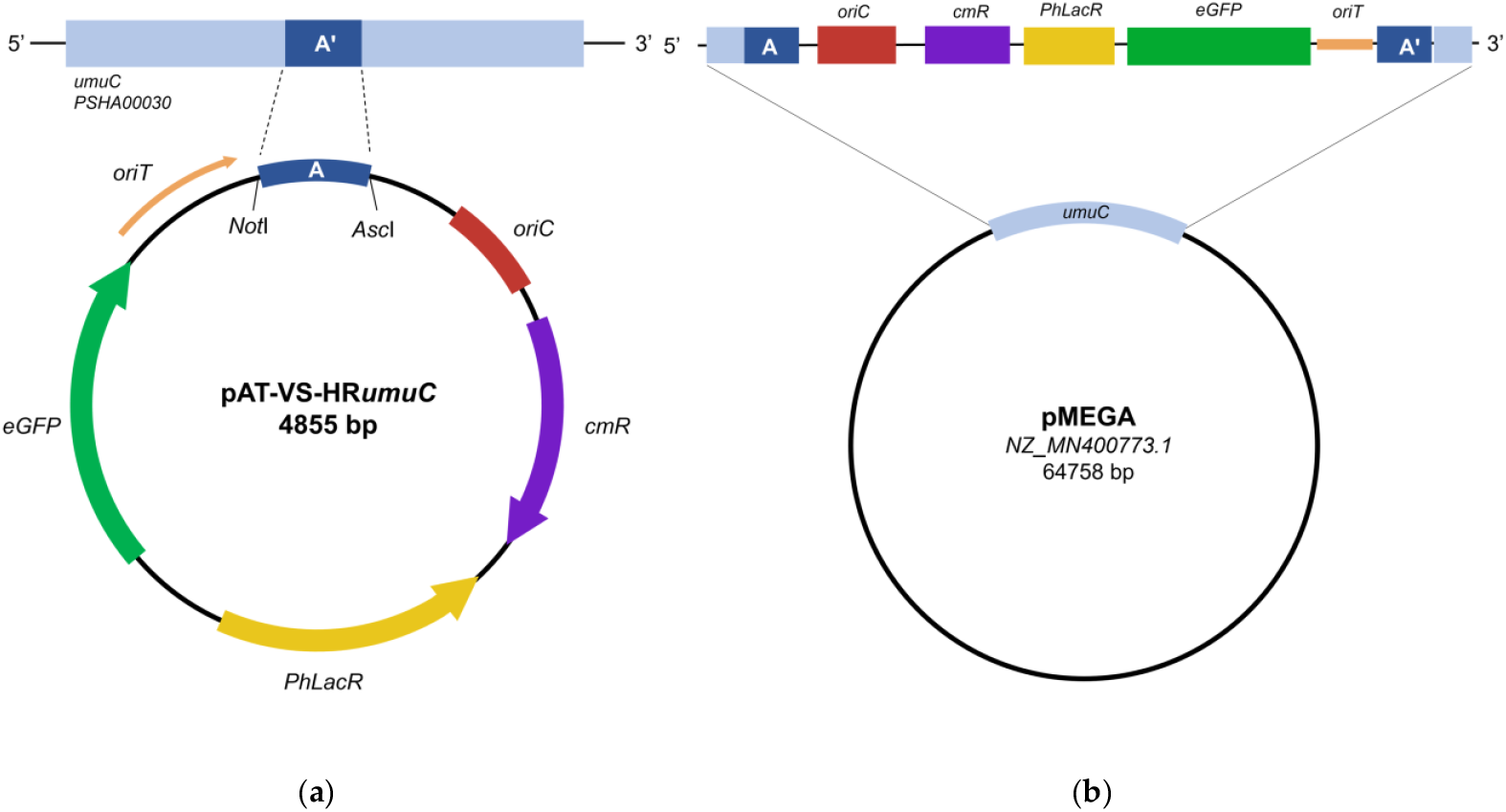
Insertion of the selection marker in pMEGA. a) Schematic representation of the suicide vector. The *umuC* homology (259 bp) region A (dark blue) is inserted between *Not*I and *Asc*I restriction site; *oriC* is the origin of replication used to propagate the plasmid in *E. coli* strains (in red); *oriT* is responsible for the initiation of the conjugative transfer (in orange); *Ph*LacR is encodes the regulator of p*LacZ* promoter (not shown); *eGFP* represents the GFP reporter under the p*LacZ* promoter (in green); *cm*R represents the chloramphenicol resistance marker (in purple). b) Representation of inserted pMEGA plasmid in KrPL insPolV mutants. The suicide vector is inserted at the *umuC* gene (in light blue) of pMEGA plasmid. The *umuC* is disrupted, gaining the cmR resistance marker and the eGFP reporter as key features.

This procedure allows the recombination event in which the pAT-VS-HR*umuC* inserts into the pMEGA plasmid in correspondence with the homology region (Figure 1b).

The suicide vector inserts at the level of the *umuC* gene encoding the catalytic subunit of the error-prone DNA polymerase V [40], both disrupting the *umu*C gene and inserting the chloramphenicol selection marker.

The presence of the eGFP was meant to be used as a selection marker to easily visualize transconjugants through fluorescence directly on agar plates. However, given the low copy number of the pMEGA plasmid, fluorescence was not easily detectable making the selection of mutants not feasible. Thus, taking advantage of the CamR selection marker, the selection of transconjugating mutants was carried out by plating cells on kanamycin, which is an episomal resistance of KrPL, and chloramphenicol TYP solid medium plates incubated at 15 °C.

Four chloramphenicol-resistant colonies were selected and detection of KrPL insPolV mutants was performed through PCR screening analyses using different couples of primers (*Nde*I_eGFP fw, *Kpn*I_eGFP rv; 5’*umuC* fw, *oriC* rv - listed in Supplementary Table S1) and bacterial DNA as a template. The PCR analysis confirmed that all four positive clones exhibit the suicide vectors inserted into the homology region definitively achieving the KrPL insPolV (Supplementary Figure S1, lanes 1-4). Subsequently, curing of pMEGA was achieved with the PTasRNA technology by designing Paired-Termini anti-sense RNA against the *repB* gene.

The antisense sequence was designed to harbour the RBS and start codon of *repB* within a loop region accessible for the targeting of the transcript mRNA as previously described by Lauro and co-workers (Supplementary Figure S2) [25]. To ensure that curing strategy did not induce any off-target effect, we performed a BLAST analysis of the *repB* antisense against *Ph*TAC125 whole genome confirming *repB* gene of pMEGA plasmid as a unique target (data not shown). The antisense mRNA sequence of *repB* (Supplementary Table S2) was cloned in the psychrophilic expression vector pB40-79-PTasRNA*lon*, an am-picillin-based derivative of pB40-79C-PTasRNA*lon* used in our previous work [25]. We exchanged the asRNA*lon* with the asRNA*repB*sequence achieving the pB40-79-PTasRNA*repB* (Supplementary Figure S3). Then, the pB40-79-PTasRNA*repB* expression vector was mobilized into the KrPL insPolV strain by intergeneric conjugation.

*3.3 Selection of the pMEGA-Cured Clones*

The expression of the PTasRNA*repB* was performed as described in the Materials and Method section 2.4, in GG 5-5 media at 15 °C and antisense production was induced using IPTG. No considerable impact on the growth behavior of the KrPL insPolV mutant was detected upon antisense production (Supplementary Figure S4). Thus, the selection of the putative cured clones and curing performance were carried out by observing the loss of chloramphenicol resistance until 32 hours after induction of the antisense-RNA on selective and non-selective TYP agar plates. The curing experiment showed that 4 clones out of about 200 clones after 32 hours from the treatment with the antisense-RNA resulted in chloramphenicol-sensitive cells. Two putative-cured clones (147, 153) were subjected to PCR analysis to prove pMEGA plasmid depletion, using bacterial DNA as a template. This screening is designed to amplify three different fragments: i) the *prom7* fragment (80 bp), which belongs to the *PSHAa_2051* gene of chromosome I (chrI) and represents a positive control of the strain, ii) the *CDS49* fragment (100 bp), a gene included in the pMEGA plasmid and iii) *umuC* (259 bp) fragment included both in pMEGA plasmid and the homology region of the inserted suicide vector. All the primers and fragments used are listed in Supplementary Table S1-S2 (prom7 fw, prom 7 rv, CDS49 fw, CDS49 rv, 5’*umuC* fw, *umuC*_*Asc*I rv). The *CDS49* and *umuC* fragments were selected to reveal the absence of the pMEGA plasmid in the cured colonies.

In a first PCR colony screening amplifying *prom7* and *CDS49*, only 2 out of 4 clones resulted to be putatively cured clones, represented by the 147 and 153 clones (data not shown). These were subjected to DNA genome extraction, and a clean PCR analysis was performed to confirm the curing of pMEGA plasmid. Indeed, the amplification of the *prom7* fragment from 147 and 153 samples, along with the absence of amplification of the *CDS49* and *umuC* confirmed the loss of the pMEGA plasmid (Supplementary Figure S5), generating two clones of a novel strain of *Pseudoalteromonas haloplanktis* TAC125 KrPL generally referred to as KrPL^2^. The applied curing procedure emphasizes the efficacy of the PTasRNA*repB* strategy to cure from a plasmid whose replication mechanism is speculated to be based on rolling circle replication (RCR) mediated by the RepB initiator protein [10]. Additionally, it undoubtedly represents a validation of the PTasRNAs conditional gene silencing technology previously designed by Lauro et al. to interfere *lon* gene expression, pointing out the versatility of the silencing system [25]. Lastly, the KrPL^2^ (147 clone) strain was subjected to culture propagation to spontaneously cure the pB40-79-PTasRNA*repB* plasmid used for pMEGA depletion.

### 3.4 Growth Performances of the Novel Strain KrPL^2^

Considering the presence of housekeeping genes harboured on pMEGA (see 3.1 section), we sought to investigate the effect of the removal of such metabolic genes on the growth patterns of KrPL^2^ strain. Thus, we focused on the evaluation of growth behaviour of the novel strain comparing its specific growth rate and cell biomass to the wt and the progenitor KrPL. The evaluation was performed at three different temperatures, 0 °C, 15 °C and 20 °C, either in the complex TYP or defined GG 5-5 media (Supplementary Figure S6).

Overall, the specific growth rate (μmax, h^-1^) of the wt strain is significantly higher if compared to its derivatives KrPL and KrPL^2^ at 15 °C and 20 °C in TYP media (Table 5).

**Table 5.** Growth kinetics parameters.

| $\mu_{\max}$ (h<sup>-1</sup>) | TYP | | | GG 5-5 | | |
| --- | --- | --- | --- | --- | --- | --- |
|  | 0 °C | 15 °C | 20 °C | 0 °C | 15 °C | 20 °C |
| wt | 0.0775 ± 0.0088 | 0.457 ± 0.023 | 0.697 ± 0.051 | 0.0345 ± 0.0035 | 0.265 ± 0.025 | 0.529 ± 0.011 |
| KRPL | 0.0567 ± 0.0048 | 0.333 ± 0.007 | 0.482 ± 0.032 | 0.0310 ± 0.0019 | 0.222 ± 0.006 | 0.318 ± 0.011 |
| KRPL<sup>2</sup> | 0.0609 ± 0.0105 | 0.327 ± 0.006 | 0.483 ± 0.019 | 0.0320 ± 0.010 | 0.230 ± 0.013 | 0.363 ± 0.015 |

| g, doubling time (h) | TYP |  |  | GG 5-5 |  |  |
| --- | --- | --- | --- | --- | --- | --- |
|  | 0 °C | 15 °C | 20 °C | 0 °C | 15 °C | 20 °C |
| wt | 9.02 ± 1.01 | 1.52 ± 0.08 | 1.00 ± 0.07 | 20.20 ± 1.99 | 2.64 ± 0.31 | 1.31 ± 0.03 |
| KRPL | 12.26 ± 0.99 | 2.08 ± 0.04 | 1.44 ± 0.09 | 22.42 ± 1.35 | 3.13 ± 0.09 | 1.85 ± 0.26 |
| KRPL<sup>2</sup> | 11.61 ± 1.93 | 2.12 ± 0.04 | 1.44 ± 0.06 | 23.74 ± 7.90 | 3.02 ± 0.16 | 1.91 ± 0.08 |

| Maximum biomass concentration (gdcw/L) | TYP |  |  | GG 5-5 |  |  |
| --- | --- | --- | --- | --- | --- | --- |
|  | 0 °C | 15 °C | 20 °C | 0 °C | 15 °C | 20 °C |
| wt | 14.22 ± 0.30 | 15.45 ± 1.91 | 12.01 ± 1.21 | 5.31 ± 0.27 | 4.73 ± 0.1 | 3.95 ± 0.02 |
| KRPL | 12.98 ± 0.19 | 10.81 ± 0.46 | 11.15 ± 0.28 | 4.86 ± 0.14 | 4.09 ± 0.23 | 3.93 ± 0.33 |
| KRPL<sup>2</sup> | 13.68 ± 0.36 | 11.24 ± 1.04 | 11.33 ± 0.59 | 4.59 ± 0.29 | 3.46 ± 0.12 | 3.95 ± 0.28 |

Instead, the specific growth rate in GG at 20 °C of KrPL^2^ is 0.363 ± 0.015 h^-1^, which is slightly higher than the 0.318 ± 0.011 h^-1^ of the KrPL strain (1,14-fold) which might indicate a likely advantage of the strain without pMEGA plasmid under this specific conditions (Table 5). Significant differences are not observed in the other tested conditions.

Accordingly, the doubling time reflects the above findings where the KrPL and KrPL^2^ show moderately higher doubling time compared to the wt cells under most of the conditions. This indicates that elimination of pMEGA in KrPL^2^ has not dramatically affected its specific growth rate.

Furthermore, in terms of biomass per liter (g_cdw_/L) no evident differences are found except in GG 5-5 at 15 °C. Compared to the wt, KrPL and KrPL^2^ yield a lower biomass concentration per liter, in which the latter reached the lowest one (Table 5). The complex TYP medium supports higher growth rates and biomass accumulation compared to the defined GG 5-5 medium, underscoring the importance of nutrient availability. Overall, no significant differences are observed in terms of growth behaviour in the specific tested medium and temperature conditions. Loss of pMEGA plasmid did not affect KrPL^2^ strains viability preserving similar growth behavior in respect to the progenitor. What stands out the most is that KrPL and KrPL^2^ are different from the wt in each tested conditions calling for further evaluation of these differences.

### 3.5 Evaluation of Plasmid Segregation Incompatibility

pMEGA is a *rep*-based plasmid in which rolling circle plasmid replication (RCR) is controlled by RepB replication initiator protein [11]. Usually, plasmid incompatibility, which emerges when plasmids share one or more elements of plasmid replication, might arise with other *rep*-based plasmids such as pKT240 [37] and pTAUp plasmids [44]. The former is a broad host range plasmid deriving from the RSF1010 plasmid belonging to the IncQ incompatibility group [34], and the latter is an endogenous rolling circle plasmid of *Psychrobacter sp*. TA144 [41]. Here we evaluated if pMEGA replication machinery is responsible for plasmid incompatibility with these two plasmids which was previously observed in the wt and KrPL strains (data not published).

The strains chosen for this analysis are the cured *Ph*TAC125 KrPL^2^, the progenitor *Ph*TAC125 KrPL, and *Ph*TAC125 wt. Each was conjugated with 3 different ampicillin-based resistance vectors: i) the broad host range pKT240 plasmid (12,392 bp), ii) pUC-oriT-pTAUp (4110 bp), iii) the psychrophilic pMAV (5009 bp). The latter is conceived as a positive control of the intergeneric conjugation experiment. Also, *Psychrobacter sp*. TAD1 was used as a positive control for intergeneric conjugation of the pUC-oriT-pTAUp, which is derived from *Psychrobacter sp*. TA144 and it is expected to have a different replication mechanism [41].

The selection of putative trans-conjugative colonies was performed at the temperature of 4 °C with the ampicillin antibiotic. However, trans-conjugative cells showed colony formation at very different times. After antibiotic selection, the cells harbouring either pKT240, pMAV or pUC18-oriT-pTAUp were replica plated on TYP agar supplemented with ampicillin. Then, clones were inoculated in TYP medium supplemented with the antibiotic. Finally, the chosen clones were subjected to plasmid extraction and DNA was analyzed through agarose gel electrophoresis but none of the examined strains harboured the *rep*-dependent plasmids. On the other hand, the positive control, pMAV, was successfully conjugated in the three strains, as well as the pUC-oriT-plasmid in the positive control *Psychrobacter sp*. TAD1. Altogether, the results suggest that broad host range pKT240 and pUC18-oriT-pTAUp plasmids were not able to replicate in *Ph*TAC125 wt, KrPL and KrPL^2^ proving that plasmid incompatibility persists, and it is not related to *repB* plasmid replication of pMEGA.

### 3.6 Evaluation of Plasmid Segregation Stability

Depletion of the pMEGA plasmid might affect the stability of heterologous introduced plasmid (pMtBL-derived). We evaluated the possible stability variability when introduced in the KrPL^2^ strain. This study compared the loss rate of the high copy number psychrophilic plasmid pB40 [42] for over 150 generations in the presence and absence of pMEGA in non-selective conditions. We compared five strains: KrPL^2^ pB40-79-PTasRNA*repB* clone 147 and clone 153, the intermediate strain KrPL insPolV pB40-79-PTasRNA*repB*, the progenitor KrPL pB40-79-PTasRNA*repB*.

Also, control of the experiment was set up exploiting the KrPL strain harbouring the pMAI-79-*eGFP* low-copy number plasmid whose stability undergoes evident reduction over time in the percentage of recombinant cells (data not published). The results show that the removal of pMEGA did not interfere with the stability of the pB40 high-copy number plasmids in the KrPL^2^ cured strains (both clones 147 and 153) and the KrPL insPolV intermediate strain (Supplementary Figure S7). The high copy number plasmids show higher stability for over 150 generations (about 100%) in each examined strain compared to the low copy number plasmid whose stability decreased from 90% after 79 generations to 65% until 150 generations (Supplementary Figure S7). The elimination of the transposase genes located in the pMEGA plasmid did not affect the stability of introduced heterologous pB40 plasmids. Also, the deletion of the *umuC* gene (inactivation of DNA polymerase V) in the intermediate strain is not involved in plasmid segregational stability.

3.7 Assessment of Plasmid Copy Number

Similarly, we evaluated if plasmid copy number (PCN) of introduced heterologous plasmids was affected. Quantitative PCR (qPCR) was performed to determine the PCN of pMAV, a psychrophilic low-copy number plasmid, comparing KrPL^2^ to the wt and KrPL strains. We evaluated the PCN in both TYP and GG 5-5 media during the exponential growth phase (Table 6). The plasmid copy number estimated in KrPL^2^ is 4.04 ± 0.34 and 6.03 ± 0.53 in TYP and GG 5-5 conditions, respectively. However, the results did not show any differences compared to the wt and KrPL strain in the GG 5-5 medium in which the estimated values are 4.68 ± 0.71 and 5.33 ± 0.69, respectively. Surprisingly, pMAV PCN in KrPL in TYP medium results in 7.99 ± 0.67 which is 2-fold and 4-fold the PCN respectively of KrPL^2^ and wt (2.61 ± 0.52).

**Table 6.** Plasmid copy number assay of pMAV plasmid.

| Strains | TYP | GG 5-5 |
| --- | --- | --- |
| wt | $2.61 \pm 0.52$ | $4.68 \pm 0.71$ |
| KrPL | $7.99 \pm 0.67$ | $5.33 \pm 0.69$ |
| KrPL<sup>2</sup> | $4.04 \pm 0.34$ | $6.03 \pm 0.53$ |

The unique behaviour of KrPL in TYP medium might be related to host cell factors, but further investigations are needed to understand this variability among the three strains. Overall, the plasmid copy number of heterologous plasmids was not affected by removal of pMEGA plasmid in the KrPL^2^ strain.

### 3.8 Evaluation of Cellular Oxidative Stress and Motility

To further explore the phenotypes conferred by the elimination of pMEGA, we focused on the ability of KrPL^2^ to cope with cellular stress evaluating its response to oxidative stress and swarming motility.

The sensitivity to oxidative agents was measured by disk-diffusion assay using 265 mM hydrogen peroxide comparing KrPL^2^, along with KrPL and wt cells upon reaching the stationary phase. After 24h of incubation at 15 °C, the growth of KrPL^2^ shows a significantly lower diameter of inhibition suggesting its higher capacity to respond to oxidative stress induced by H_2_O_2_ (Table 7).

**Table 7.**
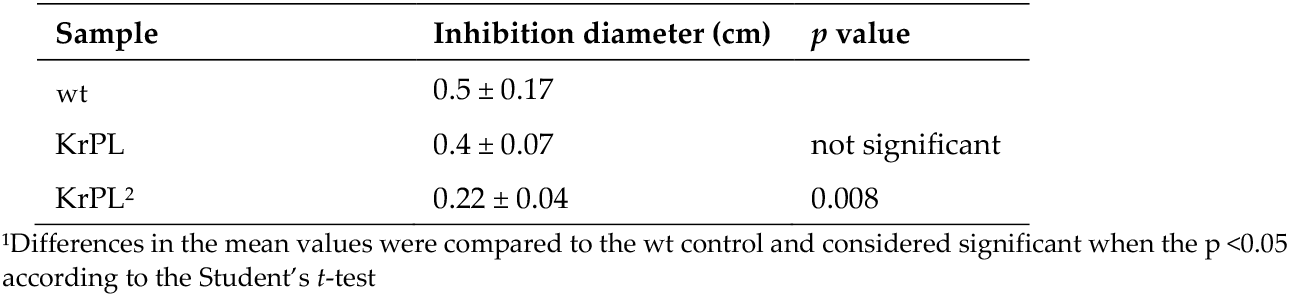
Oxidative stress assay.

Swarming ability was evaluated through a TYP soft agar motility assay and parameters were expressed as the length of the path travelled on the TYP soft-agar plates per unit of time. The bacterium KrPL^2^ displayed the same ability to move in a semi-solid TYP medium at 15 °C compared to the wt and KrPL (Supplementary Table S5).

### 3.9 Biofilm Formation Kinetics

To investigate differences in the capability of biofilm production of *Ph*TAC125 KrPL^2^ compared to the KrPL and the wt strains, cells were grown in different media at 15 °C in static conditions. Bacteria were grown in TYP and GG 5-5 at 15 °C and 0 °C in static conditions, and the biofilm was evaluated at different incubation times. The inoculum was performed at 0.2 OD/mL in 20 mL TYP and GG 5-5 supplemented with Schatz salts in 24-well plates.

The kinetics of biofilm formation were carried out for 24-, 48-, 72-, 96-, 120- and 144-hours. As shown in Figure 2, the kinetic of the biofilm formation shows a different pattern in the KrPL^2^, wt, and KrPL at 15 °C in GG 5-5. Indeed, both KrPL and KrPL^2^ were not able to accumulate biofilm when grown in GG 5-5 medium at 15 °C after 24 hours compared to the wt. But while KrPL biofilm accumulation gradually decreased over time with a lower peak of accumulation at 144 hours, KrPL^2^ shows a reduced ability to produce biofilm already after 48 hours until 144 hours. KrPL and KrPL^2^ showed differences in the production of biofilm also in GG 5-5 medium at 0 °C after 24 and 48 hours. However, after 48 hours differences between wt and KrPL and KrPL^2^ are not evident.

**Figure 2.**
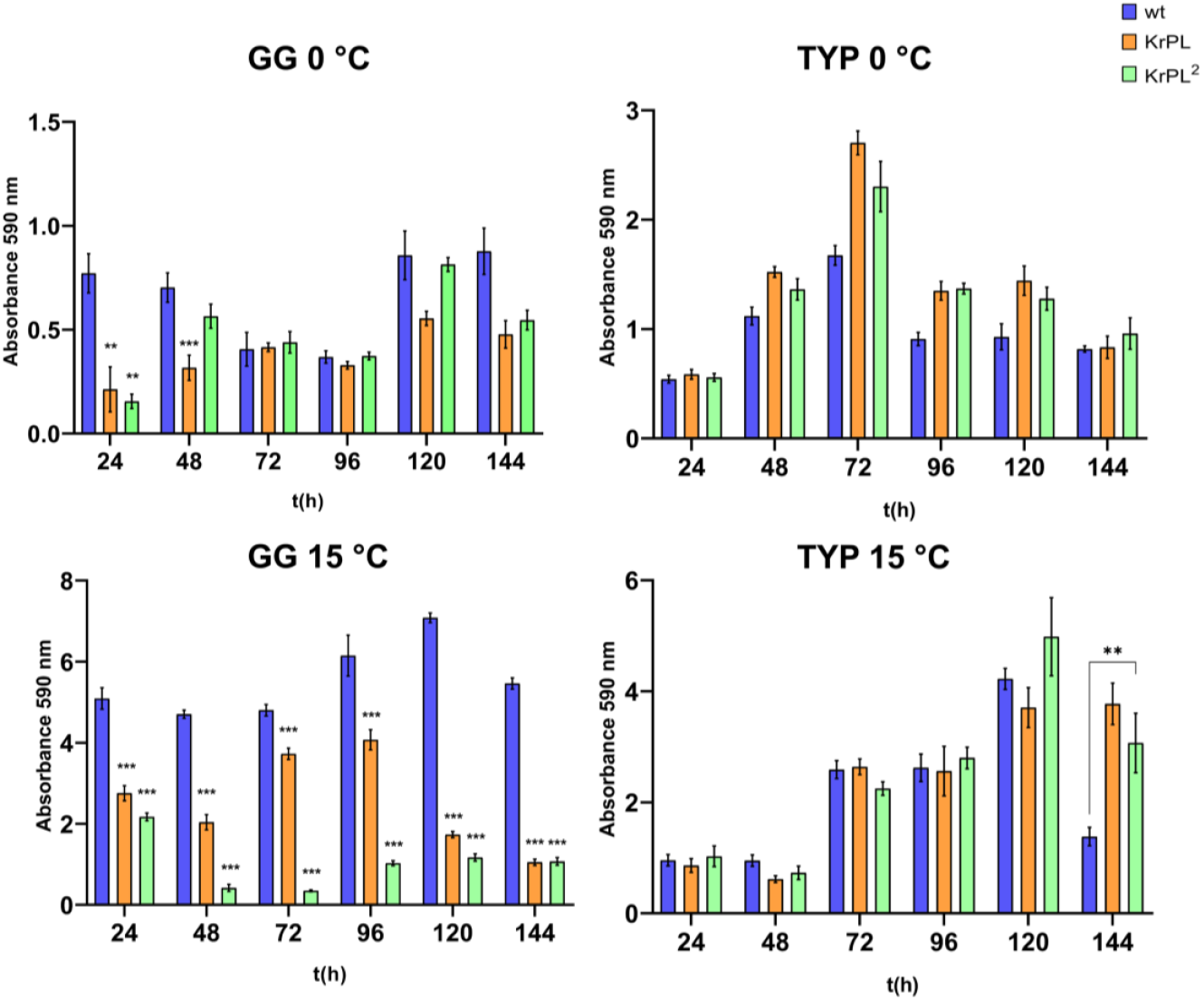
Analysis of the effect of growth medium on the *Ph*TAC125 KrPL^2^ biofilm formation at different times. Comparison of *Ph*TAC125 wt, KrPL and KrPL^2^ biofilm obtained at 15 °C and 0 °C in TYP medium and GG 5-5 medium. The biofilms were analyzed at 24 h, 48 h, 72 h, 96, 120, and 144 h with the crystal violet assay. Each data point was collected from four independent observations. Differences in the mean of absorbance were compared to the wt strain samples according to the Student’s *t*-test in which p<0.05 is considered significant (*p<0.05; **p<0.01; ***p<0.001).

Instead, in the TYP complex medium significant differences between the strains were not observed, except for the the wt that accumulated less biofilm compared to the other strains at 15 °C after 144 hours. Nevertheless, as previously described in *Ph*TAC125 wt, biofilms of KrPL and KrPL^2^ mainly accumulate at the air-liquid interface, forming pellicles at 15 °C in both BHI and GG 5-5 media [43]. Conversely, at 0 °C *Ph*TAC125 biofilm mainly accumulates at the solid-liquid interface and KrPL^2^ and KrPL begin to accumulate biofilm with 24 hours delay compared to the wt until 96 hours.

## 4. Discussion

Since *Pseudoalteromonas* is reported as a prevalent genus in marine ecosystems [44], studying its multipartite genome, particularly the presence of cryptic and mega-plasmids [29], may be of relevance for understanding its adaptation mechanisms to cold environments. In this regard, *Pseudoalteromonas haloplanktis* TAC125 represents a model microorganism for studying environmental adaptation mechanisms. Moreover, from a biotechnological perspective, the presence of the pMEGA plasmid may limiting the potential of *Ph*TAC125 as a cell-factory for difficult proteins.

In this study, genomic analyses reveal that genes encoding partitioning systems (*parA* and *parB*) and replication machinery (*repB*) are highly conserved across this genus, with over 90% sequence identity. The ParABS partitioning system is a tripartite mechanism comprising the ATPase protein ParA, the CTPase/DNA-binding protein ParB, and a centromere-like *parS* site. During cell division, ParB binds to the *parS* site, recruiting additional ParB molecules to form a protein-DNA complex, which activates the ATPase activity of ParA, driving the segregation of plasmids to opposite poles of the cell [45]. These systems, which rely on the active transfer of genetic material, are essential for the stability and retention of plasmids [46].

Transcriptomic analyses further support the stability of the pMEGA plasmid, revealing high transcriptional activity of *parAB* partitioning genes, with notable upregulation at 15 °C. This suggests that the efficient segregation and partitioning mechanism might be attributed to the ParAB system encoded by the *PSHA_00001* and *PSHA_00002* genes. Additionally, we might speculate an addition mode between pMEGA plasmid and the toxin-antitoxin genes. TA modules, which are highly transcribed at 15 °C in *Ph*TAC125, are renowned to be responsible for plasmid persistence through a mechanism called “post-segregational killing of plasmid-free segregants” that prevents plasmid-loss [50]. In an addition mode, *par* and TA module genes, might be accountable for the high inherent stability of pMEGA plasmid. However, further studies are necessary to address the putative addiction mode and prove insight into megaplasmids role in psychrophilic bacteria.

To overcome the inherent stability of the endogenous pMEGA plasmid and successfully cure it from *Ph*TAC125, we applied a dual-step approach. This involved genetic manipulation of the pMEGA plasmid to insert a chloramphenicol resistance marker through homologous recombination, followed by interference with plasmid replication using Paired-Termini antisense RNA (PTasRNA) technology. To streamline the curing procedure, we identified the *repB* gene (*PSHA_00052*) as a critical candidate, given its essential role in the plasmid replication machinery. After integrating the chloramphenicol resistance marker to facilitate the detection of plasmid loss, we proceeded to silence the *repB* gene using PTasRNA technology. This method has been shown to efficiently downregulate chromosomal genes in *Ph*TAC125, achieving high levels of silencing [25]. Our strategy proved successful in curing the pMEGA plasmid, with 4 out of 200 screened colonies identified as mutants achieving the novel KrPL^2^ strain. This curing system validates the capability of the previously developed asRNA technology in *Ph*TAC125 to silence plasmid genes and to be applied for curing of megaplasmids. Yet, we observed relatively low curing efficiency, which may be attributed to the high level of transcripts of the *repB* gene. As reported by Lauro et al., complete silencing of the *lon* gene which is highly transcribed was not observed whereas achieving complete silencing of the *PhhbO* gene, which is transcribed about 40 times less than *lon* [25]. The *repB* gene is transcribed only 3 times less compared to the *lon* gene supporting the hypothesis that high levels of transcripts may hinder gene silencing efficacy.

The KrPL^2^ strain exhibited a comparable specific growth rate to its progenitor, KrPL, across various temperatures and media, indicating that many genes encoded by the pMEGA plasmid are not essential under laboratory conditions. Its housekeeping genes, involved in replication (*repB*), partitioning (*parA* and *parB*) and plasmid maintenance (toxin-antitoxin systems), are responsible for its own replication and persistence in cells without any effect on cell viability and without imposing any metabolic burden. Although seemingly non-essential in controlled environments, pMEGA might provide benefits in environmental settings where selective pressures such as resource unavailability, competition, or environmental stressors are predominant [47]. In the context of biotechnological application of *Ph*TAC125 as a cell-factory, potential differences might be observed where further studies will highlight the importance of the KrPL^2^ strain in recombinant protein production efficiency and productivity.

Interestingly, KrPL^2^ showed enhanced tolerance to oxidative stress compared to both the wt and KrPL strains (Table 7) and biofilm formation was notably reduced in GG 5-5 medium at 15 °C decreasing overtime (Figure 2). Sequence similarity searched using pMEGA, highlighted one gene which likely might be involved in the regulation of cellular stress, the TetR/AcrR transcriptional regulator (*PSHA_p00036*). This family of transcriptional factors acts as a repressor of transcription regulating a wide range of cellular activities, including osmotic stress, and homeostasis, and thus defined as players of the global cellular stress [48]. *acr* genes also play a role in biofilm formation, such as deletion of *acrB* in *S. Typhimurium* results in impaired biofilm formation [49]. Conversely, in *A. nosocomialis acrR* deletion causes an increase in biofilm/pellicle formation [50]. However, the *Ph*TAC125 genome contains at least 10 putative *acr* genes, as reported by the MaGe Platform (MicroScope online resource), suggesting that Acr efflux pumps are part of a complex regulatory network involved in multiple bacterial functions. Additionally, many genes on pMEGA (19 of the 49 genes) are still lacking a clear functional annotation, thus it can be hypothesized that other genes have a role in the regulation of oxidative stress response and biofilm formation and it is not possible to attribute these phenotypic changes solely to the AcrR transcriptional regulator (*PSHA_p00036*).

Further studies aiming to explore the mechanisms behind these pMEGA functions are of global interest. Phenomena like controlled stress response might be interconnected with environmental stressors that these organisms face in Antarctic habitats. Their understanding will elucidate the adaptations mechanism that influence microbial survival, community dynamics and interactions. Furthermore, it is worthwhile mentioning that strains with enhanced tolerance to oxidative stress are highly appealing for industrial biotechnology. In industrial large-scale fermentation systems, oxidative stress can limit cell viability and productivity. Developing strains with improved tolerance has led to more robust and reliable production platforms as demonstrated in *C. tyrobutyricum* and *S. cerevisiae* [51,52]. In the context of reduced tendency for biofilm formation they could simplify bioreactor maintenance, minimize contamination risks, and improve overall process efficiency by reducing clogging or fouling of equipment [53].

On a conclusive note, the set up of a robust genetic modification protocol, as demonstrated by the removal of pMEGA and the achievement of the cured strain, may represent a significant achievement for the understanding of physiological role of megaplasmids in Antarctic bacteria and highlights potential biotechnological applications of *Ph*TAC125. Additionally, this study opens up the possibility to harness some of pMEGA’s valuable genetic elements, such as the origin of replication and/or promoters to repurpose for the construction of advanced genetic tools shedding light on the exploitation of microorganisms from polar ecosystems for biotechnological purposes.

## Supplementary Materials

The following supporting information can be downloaded at: www.mdpi.com/xxx/s1, Figure S1: PCR amplification of the e-GFP gene.; Figure S2: Schematic representation of the *repB* Paired Termini antisense RNA.; Figure S3: Schematic representation of PTasRNA vector.; Figure S4: Plasmid curing growth curves of *Ph*TAC125 KrPL insDNApolV strain after induction of the *repB* antisense RNA.; Figure S5: PCR screening of bacterial DNA to detect pMEGA loss.; Figure S6: Growth curves of wt, KrPL and KrPL^2^ strains in TYP and GG media at 15 °C, 0 °C and 20 °C.; Table S1: List of primers.; Table S2: List of targets.; Table S3: Motility assay of

*Ph*TAC125 strains.

## Author Contributions

A.S., C.L., A.C. and M.L.T designed the study; A.S. and C.L. developed the plasmid curing methodology; A.S. led the investigation and collected the data; A.S. conducted the data and formal analysis; A.S. wrote the original manuscript; A.S., C.L., M.C., C.R., A.C., M.F., E.P., and M.L.T. to reviewed and edited the final version; M.L.T. supervised the research activity; M.L.T. led the project administration and funding acquisition. All authors have read and agreed to the published version of the manuscript.

## Funding

This research was partially funded by the Italian “Programma Nazionale di Ricerca in Antartide” (PNRA project number PNRA18_00335) and by the National Center 5 “National Biodiversity Future Center” (code CN00000033), “Biodiversità”, financed by (PNRR)-Missione 4, Componente 2 “Dalla Ricerca all’Impresa” Investimento 1.4 “Potenziamento strutture di ricerca e creazione di campioni nazionali di R&S su alcune Key Enabling Technologies” UE—Next Generation EU, which funded the salary of Concetta Lauro.

## Data Availability Statement

The data presented in this work are available upon request from the corresponding author.

## Acknowledgments

We are grateful to the kind support by the Italian Parents’ Association “La fabbrica dei sogni 2 - New developments for Rett syndrome”.

## Conflicts of Interest

“The authors declare no conflicts of interest.”

## Notes

### Competing Interest Statement

The authors have declared no competing interest.

