## Supplementary Figures and Tables for "Engineering the Marine *Pseudoalteromonas haloplanktis* TAC125 via pMEGA Plasmid Targeted Curing Using PTasRNA Technology"

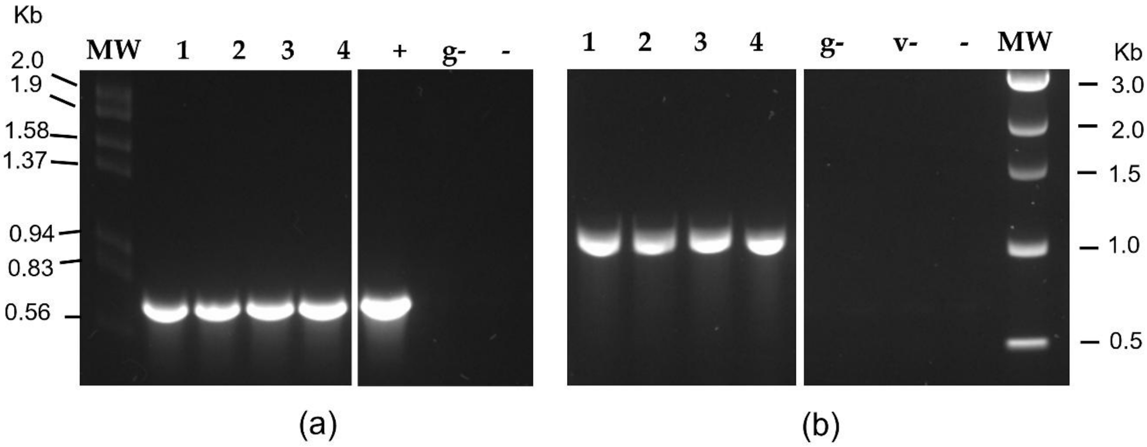


**Figure S1**. a) PCR amplification of the e-GFP gene. **M**= Lambda DNA *EcoR*I/*Hind*III ladder (Thermo Scientific); lane 1-4 represents the isolated mutant clones of KrPL pAT-VS-HR *umu*C; += pAT-VS-HR*umu*C vector as positive control g-= KrPL bacterial DNA; b) PCR analysis of the suicide vector insertion site. M= 1 kb DNA ladder (NEB); v-= pAT-VS-HR*umu*C vector negative control. Negative controls (**-**) performed without DNA template were set up for both PCR analyses. Lanes between 4 and + were removed for clarity, and the thin white line indicates the splice.

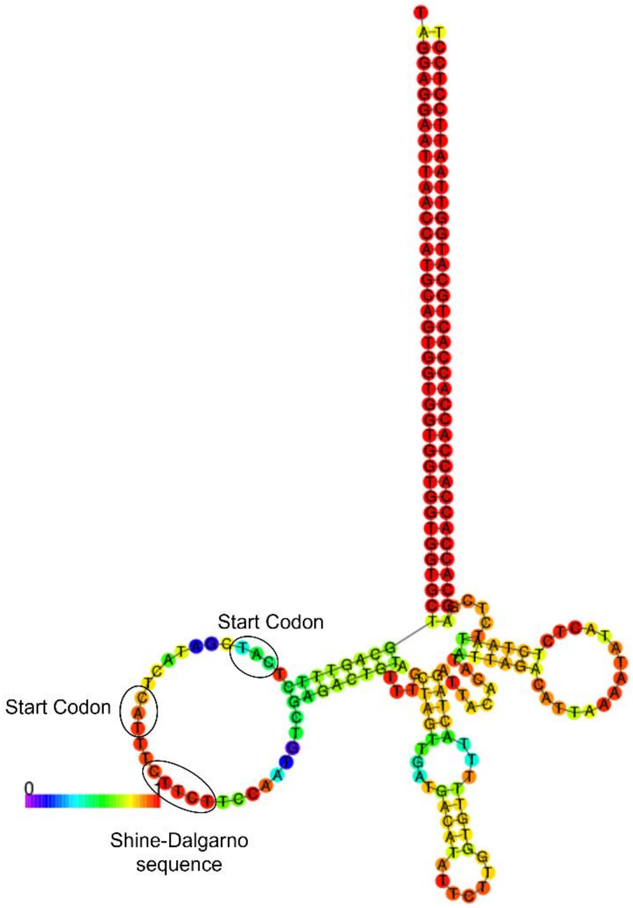


**Figure S2.** Schematic representation of the *repB* Paired Termini antisense RNA. The predicted secondary structure of the PTasRNA*rep*B through mFold tool. Start codons of translation and putative Shine-Dalgarno sequences are pointed out by circle. For the PTasRNA*rep*B two possible translation initiation sites are predicted.

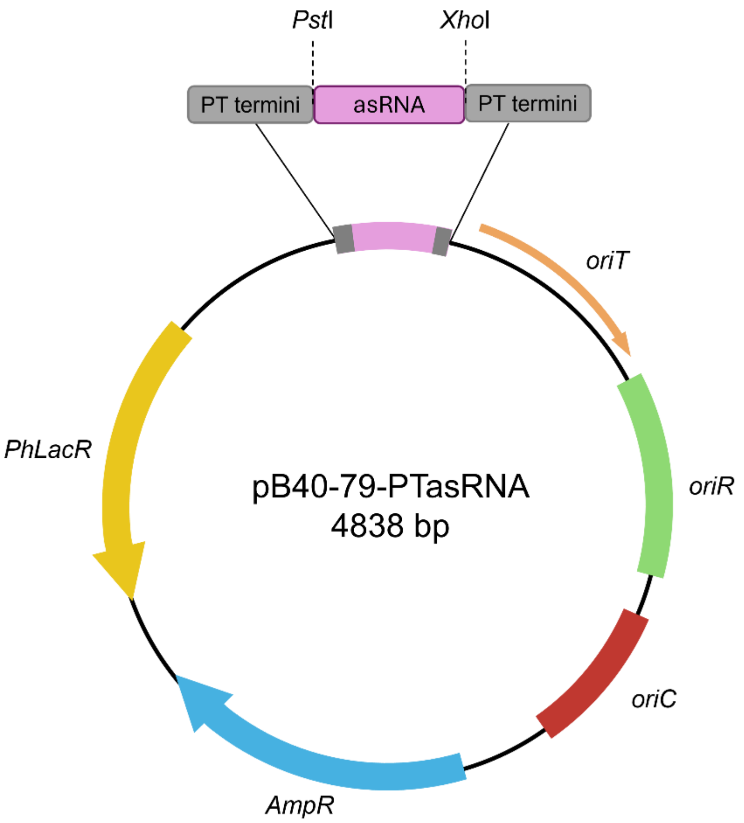


**Figure S3.** Schematic representation of PTasRNA vector. In the figure are highlighted the *Pst*I and *Xho*I restriction sites designed to insert the antisense sequence (in pink) between the two paired termini sequences (in gray). *oriC* is the origin of replication used to propagate the plasmid in *E. coli* strains (in red); *oriT* is responsible for the initiation of the conjugative transfer (in orange); *oriR* is the psychrophilic origin of replication used to propagate the plasmid in *Ph*TAC125 (in green); *Ph*LacR is encodes the regulator of p*LacZ* promoter (not shown); *amp*R represents the ampicillin resistance marker (in blue).

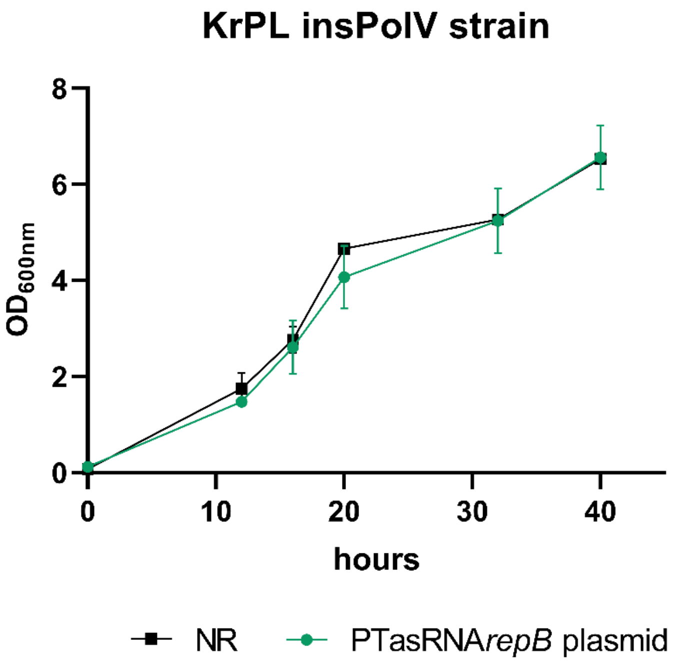


**Figure S4.** Plasmid curing growth curves of *Ph*TAC125 KrPL insDNApolV strain after induction of the *repB* antisense RNA. Growth curves of the strain expressing the PTasRNA*repB* (green) and the non-recombinant strain in black (NR) as control of the experiment. Bars indicate the standard deviation from the mean, on n=2 observations.

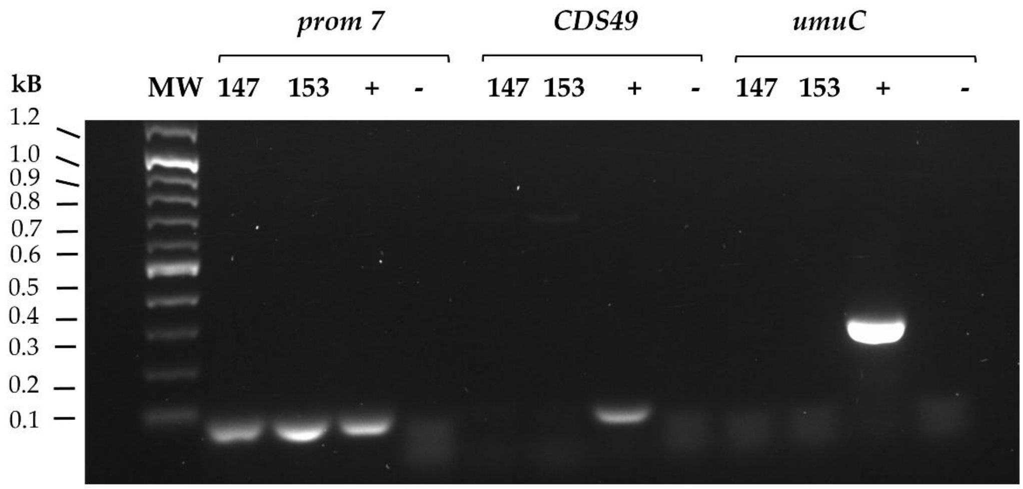


**Figure S5.** PCR screening of bacterial DNA to detect pMEGA loss. To prove deletion of pMEGA plasmid, the screening amplied three different fragments: i) the *prom7* fragment (80 bp), located on chromosome I, ii) the *CDS49* fragment (100 bp), included in the pMEGA plasmid and iii) *umuC* (259 bp) fragment included both in pMEGA and in the inserted suicide vector. *Ph*TAC125 bacterial DNA was used as template. M= Lambda DNA *EcoRI*/*HindIII* ladder (Thermo Scientific); 147 and 153: clone n°147 and n°153 selected after 32 hours induction of the PTasRNA interference plate (+) positive control using extracted genomic DNA of *Ph*TAC125 KrPL insDNApolV; (-) negative controls performed without DNA template were set up for both PCR analysis.

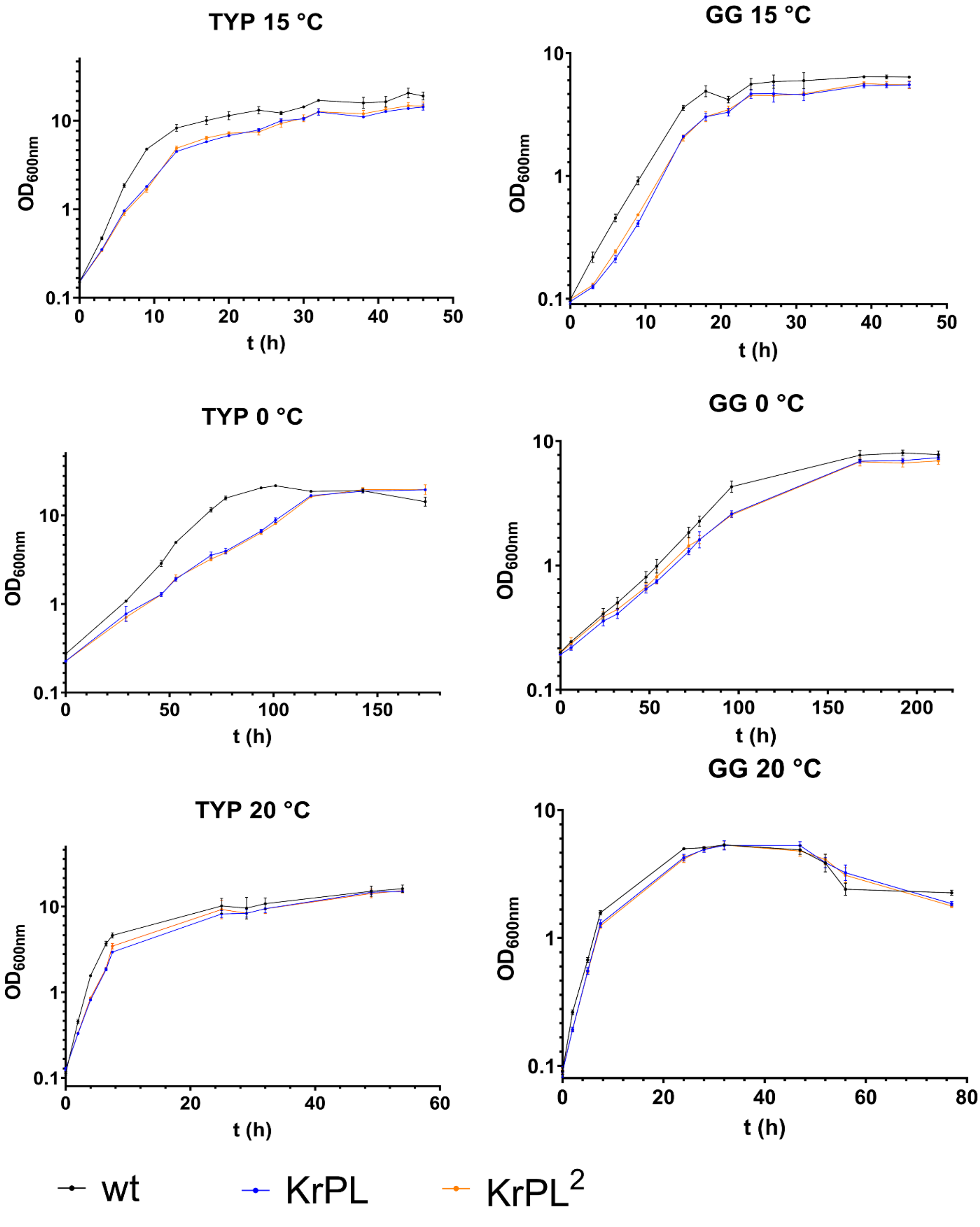


**Figure S6**. Growth curves of wt, KrPL and KrPL$^2$ strains in TYP and GG media at 15 °C, 0 °C and 20 °C

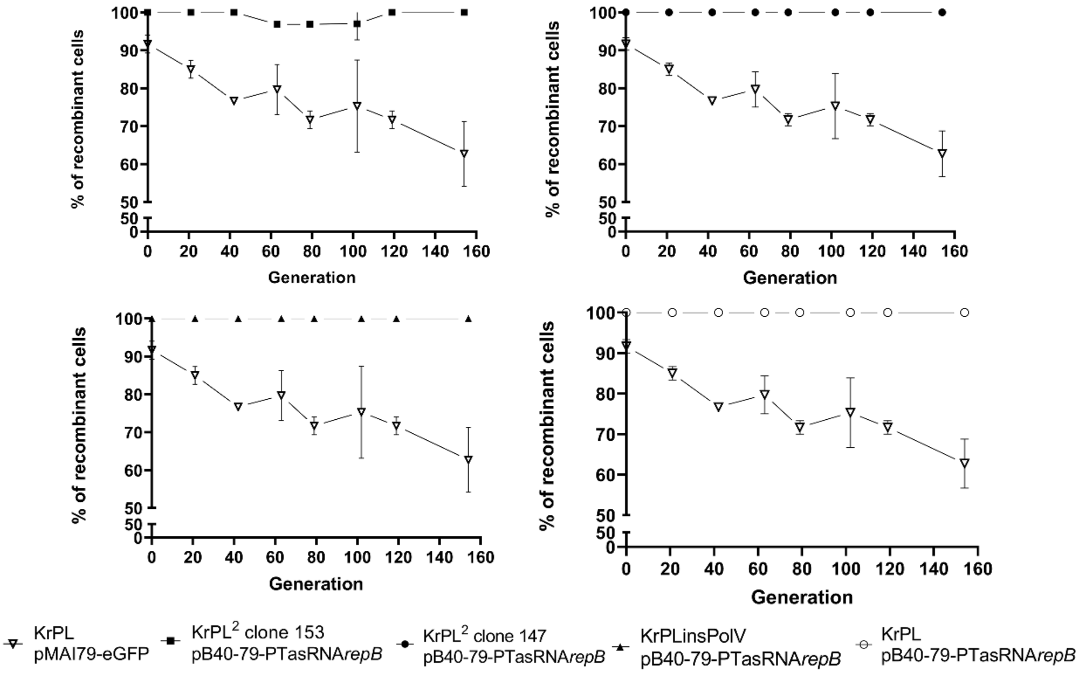


**Figure S7. Plasmid stability assay.** Plasmids pB40-79-PTasRNA*repB* in KrPL² clone 147 (●), pB40-79-PTasRNA*repB* in KrPL² clone 153 (■), pB40-79-PTasRNA*repB* in KrPL insPolV (▲), pB40-79-PTasRNA*repB* in KrPL (○), pMAI79-eGFP in KrPL (∇) were propagated for 150 generations in absence of antibiotic selection. Plasmid stability was determined by replica plating on selective media and presented as a percentage of cells that retain antibiotic resistance. Each experiment was carried out as biological duplicates and the error bars represent standard deviations.

| **Primers** | **Sequence 5'—3'** |
|---|---|
| *umuC_Not*I fw | TGCGGCCGCTAATCAGCCCAGTCATTAATG |
| *umuC_Asc*I rv | AGGCGCGCCATGGATGTATTTGTGCCATAT |
| *Nde*I_eGFP fw | ctgCATATGGGTGGAGGGAATTCAAGC |
| *Kpn*I_eGFP rv | ctGGTACCTTATTTGTAGAGCTCATCCATGCC |
| 5'*umu**C*** fw | GCTTGCAGCGAATATCACGAGTCGTTTTGC |
| *oriC* rv | CAGCGTGAGCTATGAGAAAGCGCC |
| *Xho*I-asRNA*rep*B fw | ATTTCTCGAGAATTAGAGAGTATATTTTAATGTC |
| *Pst*I-asRNA*rep*B rv | ATTCTGCAGTTTCTCATCGATACTCATTTC |
| prom7 fw | CCTTTATTCAGCGTGTTGGCGAGC |
| prom7 rv | GTTATCAGGGTCGGGCGTATCGG |
| CDS49 fw | AACTGACTGTGGTGCTCTTC |
| CDS49 rv | ACTGGTCCCTATTTGTTTATGCT |
| BlaM fw | ACTACGATACGGGAGGGCTTAC |
| BlaM rv | CGGCTGGCTGGTTTATTGCT |

**Table S1**. List of primers. The restriction enzymes sites are underlined, and the corresponding restriction enzyme are listed in the primer name

| **Target** | **Sequence 5'—3'** | **Length (bp)** |
|---|---|---|
| as*rep*B | TTTCT**CAT**CGATACT**CAT**TTCTTCTTCCAATGTCGAGACTGTATTTGCTAGTTGATGACAT ATTCTTGGTGTTTTTACTAGATTACACATATTAGACATTAAAATATACTCTCTAATT | 110 |
| prom7 | CCTTTATTCAGCGTGTTGGCGAGCACCGACACGACAGCGTACTACTTGGCTGGGCTGCCG ATACGCCCGACCCTGATAAC | 80 |
| CDS49 | AACTGACTGTGGTGCTCTTCTGAGCTAGTTGTATTTATCGCTCTATTCAGTTCACTAGCAT AAACAAATAGGGACCAGT | 79 |
| *umuC* | TAATCAGCCCAGTCATTAATGATTTTAGAAAAACCAGTAAAGCGTAAAAATGACTCATC AATACTATAAACATAATGCTCATCACAAAAACGATTAATAACCGTCATCATTCTCTCACT CAAATCAGCGTAAAGTTCATAATTTGAAGAGCGTACGATGACATTATGCTGCTCTAAATA GGGTTTAATTTTGAAGTAAGGTTCAAATTTCGGGATATTGAGCTTTCTAGCAATAGGACA TATGGCACAAATACATCCAT | 259 |

**Table S2**. List of targets. The Shine-Dalgarno sequences are double underlined, whereas the start codons are reported in bold.

| **Strain** | **24 h** | **48 h** | **72 h** |
|---|---|---|---|
| wt | 1.07 ± 0.12 | 1.37 ± 0.06 | 2.03 ± 0.25 |
| KrPL | 1.00 ± 0.01 | 1.50 ± 052 | 2.17 ± 0.50 |
| KrPL$^{2}$ | 1.07 ± 0.06 | 1.57 ± 0.4 | 2.10 ± 0.66 |

**Table S3**. Motility assay of *Ph*TAC125 strains. Swarming ability was evaluated spotting the culture on TYP soft media (0.3%) for 72 hours of incubation at 15 °C. The length of the path is calculated as cm of path travelled on the media per unit of time. The experiment was performed as three technical replicates. The error is reported as standard deviation.
